# Assessing inhibitors of mutant isocitrate dehydrogenase using a suite of pre-clinical discovery assays

**DOI:** 10.1101/124024

**Authors:** Daniel J. Urban, Natalia J. Martinez, Mindy I. Davis, Kyle R. Brimacombe, Dorian M. Cheff, Tobie D. Lee, Mark J. Henderson, Steven A. Titus, Rajan Pragani, Jason M. Rohde, Yuhong Wang, Surendra Karavadhi, Pranav Shah, Olivia W. Lee, Amy Wang, Andrew McIver, Hongchao Zheng, Xiaodong Wang, Xin Xu, Ajit Jadhav, Anton Simeonov, Min Shen, Matthew B. Boxer, Matthew D. Hall

## Abstract

Isocitrate dehydrogenase 1 and 2 (IDH1 and IDH2) are key metabolic enzymes that are mutated in a variety of cancers to confer a gain-of-function activity resulting in the accumulation and secretion of an oncometabolite, D-2-hydroxyglutarate (2-HG). Accumulation of 2-HG can result in epigenetic dysregulation and a block in cellular differentiation, suggesting these mutations play a role in neoplasia. Based on its potential as a cancer target, a number of small molecule inhibitors have been developed to specifically inhibit mutant forms of IDH (mIDH1 and mIDH2). Here, a panel of mIDH inhibitors were systematically profiled using biochemical, cell-based, and tier-one ADME techniques. We quantified the biochemical effect of each inhibitor on mIDH1 (R132H and R132C) and mIDH2 (R172Q). The effect of these inhibitors on 2-HG concentrations in seven cell lines representing five different IDH1 mutations in both 2D and 3D cell cultures was assessed. Target engagement of these inhibitors was analyzed utilizing cellular thermal shift assays (CETSA), the effects of inhibitors on reversing 2-HG-induced block on leukemic cellular differentiation. We conclude from our mIDH1 assay panel that AG-120 and a Novartis inhibitor exhibited excellent activity in all biochemical and most cellular assays. While AG-120 has superior DMPK properties, it lacks efficacy a leukemic differentiation model. In conclusion, we present a comprehensive suite of *in vitro* preclinical drug development assays that can be used as a tool-box to identify lead compounds for mIDH drug discovery programs, as well as what we believe is the most comprehensive publically available dataset on the top mIDH inhibitors.

## Introduction

The mutant isocitrate dehydrogenases 1 (mIDH1) and 2 (mIDH2) represent a remarkable example of the rapidity with which target identification can translate to small-molecule drug discovery and clinical trials (1,2). Wild type IDH1 protein forms a homodimer that catalyzes the conversion of isocitrate to α-ketoglutarate (α-KG, also termed 2-oxoglutarate, 2-OG), using the co-factor NADP^+^ (3). Studies in 2009/10 demonstrated that a subset of acute myelogenous leukemias (AML) and gliomas harbored heterozygous mutations at the R132 position of IDH1 and R140 or R172 of IDH2 (mutations in the two genes are mutually exclusive) (4-6). Subsequently, it has been shown that >75% of low-grade gliomas and 20% of AML have mutations in IDH1 or IDH2, and mutations are also found in other solid tumors such as chondrosarcoma, cholangiocarcinoma, colon, pancreatic and prostate cancer to varying extents (1,7).

While the R132 mutation reduces normal enzymatic activity, it also confers a neomorphic (gain-of-function) activity. Metabolic profiling showed that the wild type IDH1 product, α-KG, is the substrate of mIDH1, producing D-2-hydroxyglutarate (2-HG) in a NADPH-dependent manner (8). 2-HG is detected at low concentrations in normal cells, yet is significantly elevated in tumor cells (up to 10 mM) and plasma of patients bearing IDH1/2 mutations (8). Considerable evidence now exists demonstrating that the 2-HG ‘oncometabolite’ produced by mIDH1 plays a role in tumorigenesis and cellular proliferation (9-12). 2-HG is structurally similar to α-KG and has been shown to inhibit α-KG-dependent enzymes, modifying the endogenous cellular biochemical balance (13). Consequently, inhibition of α-KG-dependent histone and DNA demethylases by 2-HG leads to elevated methylation of these substrates, altered gene expression, and a block to cell differentiation (14).

Inhibition of oncogenic mIDH1/2 represents an opportunity for therapeutic intervention. As a target, mIDH1/2 contains a distinct genetic modification allowing personalized medicine using tumor gene sequencing and oncometabolite detection as biomarkers (15). Ideally, specific inhibition of mIDH1 or mIDH2 would have few clinical side-effects, as no endogenous biochemistry would be disrupted by pharmacologic modulation (analogous to the oncogene BCR-ABL). mIDH1 and mIDH2 have therefore received significant attention for the development of small molecule inhibitors (2).

Several biotech and pharmaceutical discovery campaigns for mIDH1/2 inhibitors have been disclosed. Many of these inhibitors are in clinical trials for patients with AML or solid tumors demonstrating the rapid advancement from the first report of the IDH1 mutation seven years ago to current late phase clinical trials. Agios Pharmaceuticals has reported the mIDH1 inhibitor AG-5198 (16), optimized from a phenyl-glycine hit from a biochemical screen (17), along with a similar lead, the phenyl-glycine analog ML309, in collaboration with NCATS (18). Agios has subsequently revealed their mIDH1 inhibitor clinical candidate, AG-120, currently in phase III (19). Likewise, Novartis has reported multiple probe molecules and their clinical candidate, IDH305 (20), is currently in Phase I. Similarly, Sanofi and GlaxoSmithKline have reported their own mIDH1 probe molecules (21,22). Agios has also developed mIDH2 inhibitors, derived from biochemical screens to produce the heterocyclic urea sulfonamide probe, AGI-6780 (23), and two clinical candidates, AG-221 (Phase III) and AG-881 (Phase I, pan-mutant IDH1/2 inhibitor) (20). Along with these published peer-reviewed reports, additional chemotypes have been disclosed in patents, which have led to the development of several commercially available inhibitors with limited characterization.

While multiple inhibitors have been reported, some of which are available to research laboratories, the clinical development and proprietary nature of mIDH inhibitors has meant that limited characterization is publically available. In addition, the scope of activity of many of these mIDH inhibitors as pre-clinical chemical probes or tool molecules has not been assessed, and no insight into relative utility is currently available. To remedy this situation, we comprehensively characterized the activity of nine chemically diverse mIDH1 inhibitors in a panel of *in vitro* biochemical as well as functional cellular assays. Specifically, inhibitor activity was assessed *in vitro* against mIDH1 enzymes containing multiple clinically-relevant mutations. Cell-based activity was assessed by measurement of 2-HG production, via mass-spectrometry as well as an in-house developed high-throughput fluorescence assay, in cell lines harboring somatic or engineered mIDH1. Inhibitor target engagement was confirmed by utilizing the cellular thermal shift assay (CETSA) (24). Rescue of the 2-HG induced blockade on leukemic differentiation was assessed to compare compound efficacy in an *in vitro* AML model. Three-dimensional (3D) culture of mIDH1 lines was developed as an *in vitro* solid tumor model to compare inhibitor activity via 2-HG production and cell growth. Finally, physicochemical and *in vitro* drug metabolism parameters (i.e. solubility, PAMPA permeability, Caco-2 cell transwell permeability, plasma protein binding, and liver microsomal stability) were determined. Altogether, these assays provide a comprehensive assessment of the comparative activity of the mIDH1 inhibitor landscape and a platform that could be implemented for future mIDH1 inhibitor discovery campaigns. This effort also illustrates a rigorous and general approach to addressing reproducibility in preclinical drug discovery through construction of a hierarchy of biological assays to assess compound suitability for clinical development. Reproducibility and target validity is assessed based on replication of activity not of a single compound in a single assay, but rather of many compounds tested repeatedly across multiple assays with diverse readouts (25).

## Materials and Methods

### Compounds

Novartis 224 (HY-18717), Novartis 556 (HY-13972), Agios 135 (HY-12475), and AG-221 (HY-18690) were purchased from Medchem Express. AGI-5198 (14624) and AGI-6780 (14639) were purchased from Cayman Chemicals. IDH2-C100 was purchased from XcessBio. AG-120 was purchased from CG Biopharma. GSK864 was a gift from Structure Genomics Consortium (21). ML309 was synthesized as previously described (18). The synthesis of Sanofi 1 and Novartis 530 are reported in the supplemental methods.

### IDH1 and IDH2 biochemical assays

WT IDH1 protocol was performed as previously described (18). Generally, WT IDH1/2 enzyme in assay buffer was added to the black solid bottom 1,536-well assay plate. A pintool (Kalypsys) was used to transfer 23 nL of compound solution (library and control) to the 1,536-well assay plates. After 30 min of incubation at room temperature, 1 μL of substrate buffer was added to initiate the reaction. The plate was rapidly transferred to a ViewLux (PerkinElmer) and the fluorescence product resorufin was measured in kinetic mode. Additional details are described in supplemental methods.

### mIDH1-R132H and IDH-R132C enzyme assay

mIDH1 protocol was performed as previously described (18). Generally, 3μL of mutant enzyme in assay was added to the black solid bottom 1,536-well assay plate. A pintool (Kalypsys) was used to transfer 23 nL of compound solution (library and control) to the 1,536-well assay plates. After 30 min incubation at room temperature, 3 μL of buffer containing NADPH and α-KG was added to initiate the reaction. After a 40-50 min incubation period, 3 μL of detection buffer was added to a total volume of 9 μL. The plate was transferred to an Envision (PerkinElmer) after 5 min and the fluorescence product resorufin was measured. Additional details are described in supplemental methods.

### mIDH2-R170Q enzyme assay

Reaction Biology profiled all compounds against mIDH2-R172Q using a diaphorase / resazurin coupled reaction.

### Cell culture

U87 (R132H) and HT1080 were obtained from Agios, THP1 (WT and R132H) were obtained from Ravindra Majeti (Stanford University), SNU1079 and RBE from Nabeel M. Bardeesy (Harvard University), and JJ012 were obtained from Karina Galoian (University of Miami). Short tandem repeat analysis was performed on each cell line using WiCell Research Institute and Genetic Resources Core Facility at the University Johns Hopkins (Supplemental Table 2). Additional cell culture and 2-HG production assay details are described in supplemental methods.

### Diaphorase coupled D-2-Hydroxyglutarate Dehydrogenase Assay

The diaphorase coupled D-2-hydroxyglutarate dehydrogenase assay was a modification of the report by Jorg Balss and colleagues (26). HT1080 cells (4,000 cells/well) were dispensed into 384-well TC-treated cell culture plates. A pintool (Kalypsys) was used to transfer 96 nL of compound solution to the 384-well assay plates. After 48 hr incubation at 37°C, 10 μL of media was transferred to new 384-well white solid bottom assay plates.

Collected media was neutralized and 36 μL of detection solution (containing active recombinant D2HGDH) was then added to the 12 μL of neutralized media. After 30 min incubation at room temperature, the plates were transferred to a Viewlux HTS Microplate Imager (PerkinElmer) and the fluorescence product resorufin was measured. Additional details are described in supplemental methods.

### Cellular thermal shift assays

The cellular thermal shift assay and the isothermal dose response was run as previously described (27). Additional details are described in supplemental methods.

### THP1 monocytic differentiation assay

Human THP1 leukemia cells engineered to express either wild type IDH1 or mutant IDH1-R132H under a doxycycline-inducible promoter have been previously described (28). To induce transgene expression, cells were treated with doxycycline (2 μg/mL final). After 4 days of doxycycline induction, cells were pre-treated with either vehicle DMSO or the indicated mIDH1 inhibitor at a final concentration of 0.5 μM for 4 additional days. Monocyte differentiation was induced by the addition of phorbol-12-mysistate-13-acetate (PMA) following a previously described protocol (29) with some modifications. Attached cells were washed, stained, imaged and quantified. Additional details are described in supplemental methods.

### Histone methylation assay

Human THP-1 leukemia cells were grown as previously described. Following treatment, cells (6×10^6^) were then collected, pelleted, and snap frozen. Total histone extracts were prepared using a Histone Extraction Kit (Abcam #113476). Extracts (7 μg) were separated on 4-12% Bis-Tris NuPAGE gels (Thermo Fisher) and transferred to nitrocellulose membranes using an iBlot transfer system (Thermo Fisher). Primary antibodies were diluted in TBST + 5 % BSA and incubated overnight at 4°C: mouse anti-H3K4me3 (Abcam #1012; 1:500), rabbit anti-H3K9me3 (Cell Signaling #13969; 1:1000) and rabbit anti-total H3 (Cell Signaling #4499; 1:2000). Membranes were washed 3 × 10 min with TBST and incubated with HRP-conjugated secondary antibodies (1:5000 in TBST + 1% BSA) for 1 hr at room temperature. SuperSignal West Dura chemiluminescence substrate (Thermo Fisher) was added and immunoblots were imaged using a BioRad ChemiDoc Imager. Densitometric analysis was performed using ImageJ. Additional details are described in supplemental methods.

### In Vitro Physicochemical and Drug Metabolism Studies

High-Throughput PAMPA assay, High-Throughput kinetic solubility test assay, Microsomal metabolic stability assay, Human plasma protein binding assay, Caco-2 permeability assay, Bidirectional MDCK-MDR1 permeability assay details are described in supplemental methods.

## Results

### Biochemical characterization of mIDH1inhibitors

We characterized the activity of 9 mutant IDH1 inhibitors ( Supplemental Figure 1, A-I) and 3 control mIDH2 inhibitors (Supplemental Figure 1, J-L) against purified recombinant R132H and R132C mIDH1 enzymes, as well as the purified wild type IDH1 and IDH2 enzymes. Each enzyme was coupled to a diaphorase/resazurin readout for a red-shifted fluorescence detection of cofactor turnover to avoid compound fluorescence interference (Figure 1A) (18,30,31). All mIDH1 inhibitors were more potent against both R132H and R132C mIDH1 enzymes than wild type IDH1, and were inactive against wild type IDH2 (Figure 1B-D and Table 1). The most potent inhibitors against both mIDH1 enzymes were AG-120 and Novartis 530 with IC_50_ values of 40-50 nM. Additionally, AG-120 and Novartis 530 revealed an 85 – 106 and 70 - 88 fold separation in potency, respectively, relative to wild type IDH1 (Figure 1B-D and Table 1). A strong correlation (R^2^ = 0.85) was found for cross-inhibition of the R132H and R132C enzymes (Figure 1E). Sanofi’s compound 1 demonstrated weak activity (IC_50_ >13 μM) against all enzymes, and was excluded from subsequent cell-based studies. Control mIDH2 inhibitors demonstrated weak activity (>10 μM) against mutant and wild type IDH1 enzymes, and wild type IDH2 enzyme (Supplemental Table 1).

**Figure 1.**
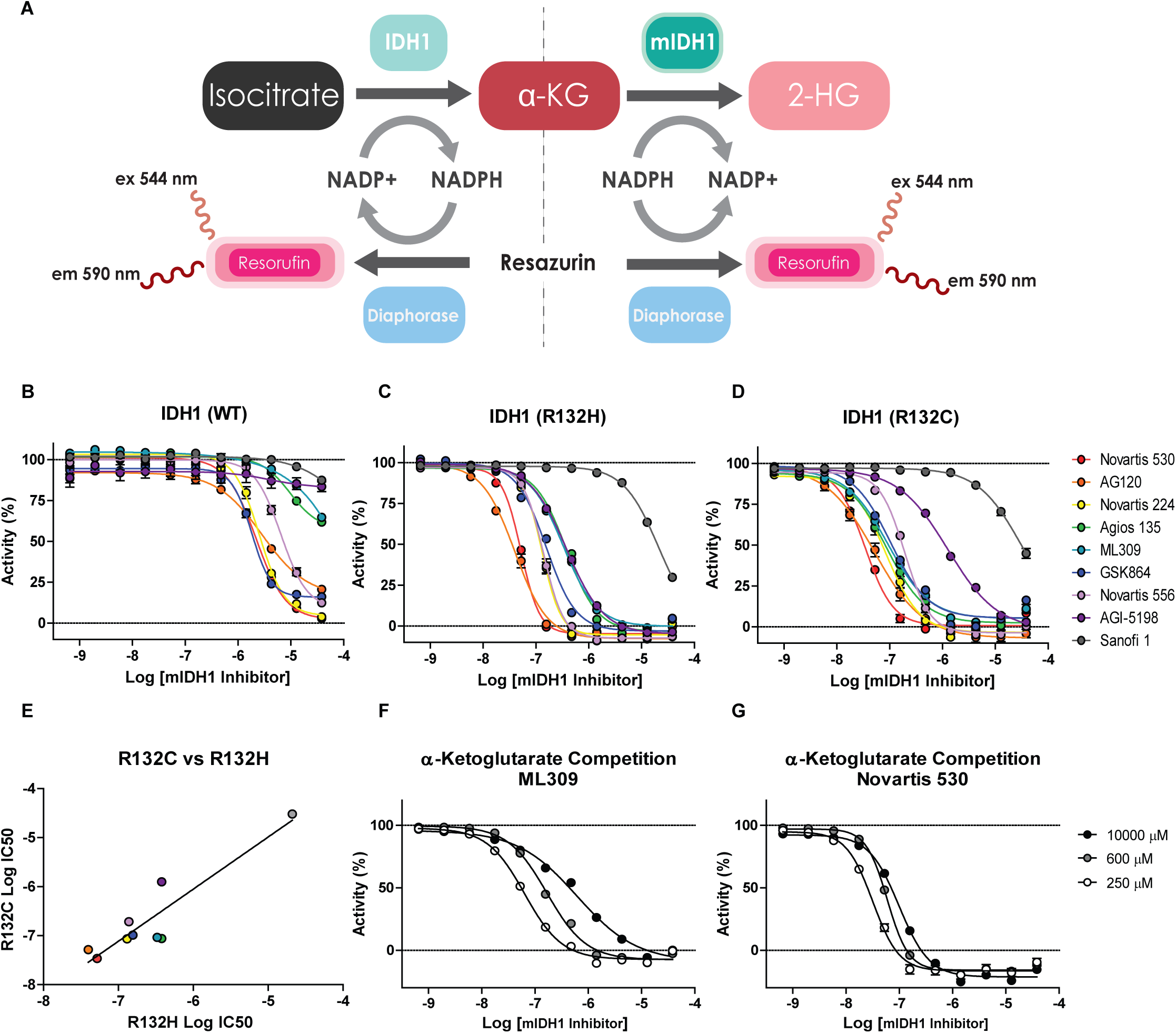
Biochemical characterization of mIDH1 inhibitors. A, Scheme of IDH1 and mIDH1 enzymatic reactions coupled to diaphorase for fluorescence detection. B-D, Dose-response analyses show the selectivity of mIDH1 inhibitors on the activity of purified (B) wild type IDH1, (C) R132H and (D) R132C mIDH1 enzymes (n=3). E, Comparison of mIDH1 inhibitors Log IC_50_ values on the activity of R132H and R132C mIDH1 enzymes show a strong correlation (R^2^ = 0.85, P value = 0.001). A large dose-response shift is shown for (F) ML309 compared to (G) Novartis 530 when using a range of concentrations of α-KG, suggesting ML309 acts as a competitive inhibitor (n=3).

**Table 1-.**
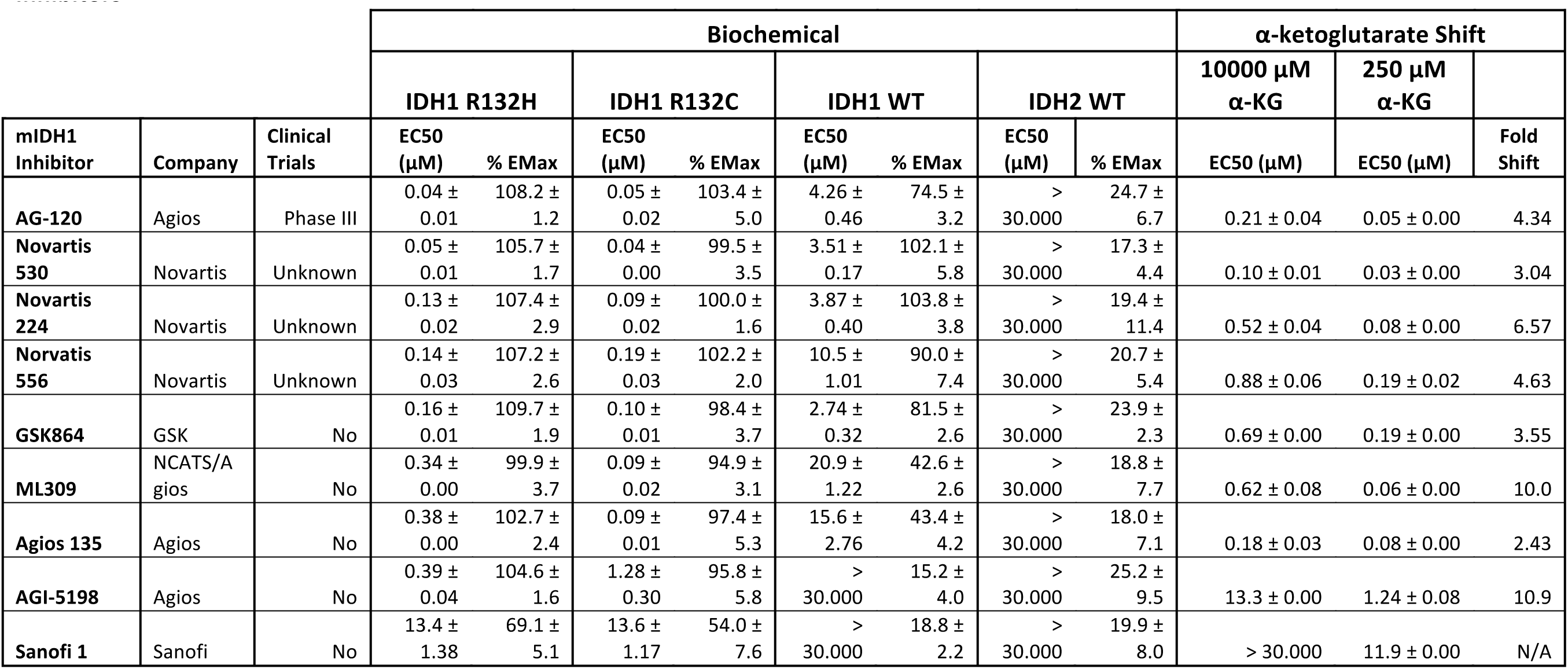
Biochemical Profile of mIDH1 Inhibitors.

| mIDH1 Inhibitor | Company | Clinical Trials | Biochemical |  |  |  |  |  |  |  | α-ketoglutarate Shift |  |  |
| --- | --- | --- | --- | --- | --- | --- | --- | --- | --- | --- | --- | --- | --- |
|  |  |  | IDH1 R132H |  | IDH1 R132C |  | IDH1 WT |  | IDH2 WT |  | 10000 μM α-KG | 250 μM α-KG |  |
|  |  |  | EC50 (μM) | % EMax | EC50 (μM) | % EMax | EC50 (μM) | % EMax | EC50 (μM) | % EMax | EC50 (μM) | EC50 (μM) | Fold Shift |
| **AG-120** | Agios | Phase III | 0.04 ± 0.01 | 108.2 ± 1.2 | 0.05 ± 0.02 | 103.4 ± 5.0 | 4.26 ± 0.46 | 74.5 ± 3.2 | > 30.000 | 24.7 ± 6.7 | 0.21 ± 0.04 | 0.05 ± 0.00 | 4.34 |
| **Novartis 530** | Novartis | Unknown | 0.05 ± 0.01 | 105.7 ± 1.7 | 0.04 ± 0.00 | 99.5 ± 3.5 | 3.51 ± 0.17 | 102.1 ± 5.8 | > 30.000 | 17.3 ± 4.4 | 0.10 ± 0.01 | 0.03 ± 0.00 | 3.04 |
| **Novartis 224** | Novartis | Unknown | 0.13 ± 0.02 | 107.4 ± 2.9 | 0.09 ± 0.02 | 100.0 ± 1.6 | 3.87 ± 0.40 | 103.8 ± 3.8 | > 30.000 | 19.4 ± 11.4 | 0.52 ± 0.04 | 0.08 ± 0.00 | 6.57 |
| **Norvatis 556** | Novartis | Unknown | 0.14 ± 0.03 | 107.2 ± 2.6 | 0.19 ± 0.03 | 102.2 ± 2.0 | 10.5 ± 1.01 | 90.0 ± 7.4 | > 30.000 | 20.7 ± 5.4 | 0.88 ± 0.06 | 0.19 ± 0.02 | 4.63 |
| **GSK864** | GSK | No | 0.16 ± 0.01 | 109.7 ± 1.9 | 0.10 ± 0.01 | 98.4 ± 3.7 | 2.74 ± 0.32 | 81.5 ± 2.6 | > 30.000 | 23.9 ± 2.3 | 0.69 ± 0.00 | 0.19 ± 0.00 | 3.55 |
| **ML309** | NCATS/Agios | No | 0.34 ± 0.00 | 99.9 ± 3.7 | 0.09 ± 0.02 | 94.9 ± 3.1 | 20.9 ± 1.22 | 42.6 ± 2.6 | > 30.000 | 18.8 ± 7.7 | 0.62 ± 0.08 | 0.06 ± 0.00 | 10.0 |
| **Agios 135** | Agios | No | 0.38 ± 0.00 | 102.7 ± 2.4 | 0.09 ± 0.01 | 97.4 ± 5.3 | 15.6 ± 2.76 | 43.4 ± 4.2 | > 30.000 | 18.0 ± 7.1 | 0.18 ± 0.03 | 0.08 ± 0.00 | 2.43 |
| **AGI-5198** | Agios | No | 0.39 ± 0.04 | 104.6 ± 1.6 | 1.28 ± 0.30 | 95.8 ± 5.8 | > 30.000 | 15.2 ± 4.0 | > 30.000 | 25.2 ± 9.5 | 13.3 ± 0.00 | 1.24 ± 0.08 | 10.9 |
| **Sanofi 1** | Sanofi | No | 13.4 ± 1.38 | 69.1 ± 5.1 | 13.6 ± 1.17 | 54.0 ± 7.6 | > 30.000 | 18.8 ± 2.2 | > 30.000 | 19.9 ± 8.0 | > 30.000 | 11.9 ± 0.00 | N/A |

Given the difficulty of predicting the physiological substrate and co-factor concentrations in target tissues and the fact that these are in constant flux within the cell, at times it can be beneficial to pursue non-competitive mechanisms of inhibition (32). To determine if any of the mIDH1 inhibitors demonstrated competitive inhibition with the mIDH1 substrate α-KG, enzyme inhibition was assessed in the presence of high (10 mM), medium (600 μM), and low (250 μM) concentrations of α-KG. ML309 and AGI-5198 revealed a >10-fold loss of activity in the presence of high α-KG, whereas the strongest inhibitors, AG-120 and Novartis 530 (Table 1), demonstrated only a 3-4 fold IC_50_ shift (Figure 1F-G).

Inhibition of R172Q mIDH2 enzyme was assessed for all compounds; with the characterized mIDH2 inhibitors demonstrating strong inhibition (Supplementary Figure 2A and Supplemental Table 1). The Agios probe AGI-6780 (IC_50_ = 23 nM) was slightly more potent than the clinical candidate, AG-221 (IC_50_ = 65 nM), and demonstrated greater inhibitory activity than the patent compound IDH2-C100 (IC_50_ ∼1 μM). None of the mIDH2 inhibitors demonstrated activity against the IDH1 enzymes, but the mIDH1 inhibitor GSK864 (mIDH1 IC_50_ = 100-150 nM) did inhibit mIDH2 (IC_50_ = 183 nM).

**Figure 2.**
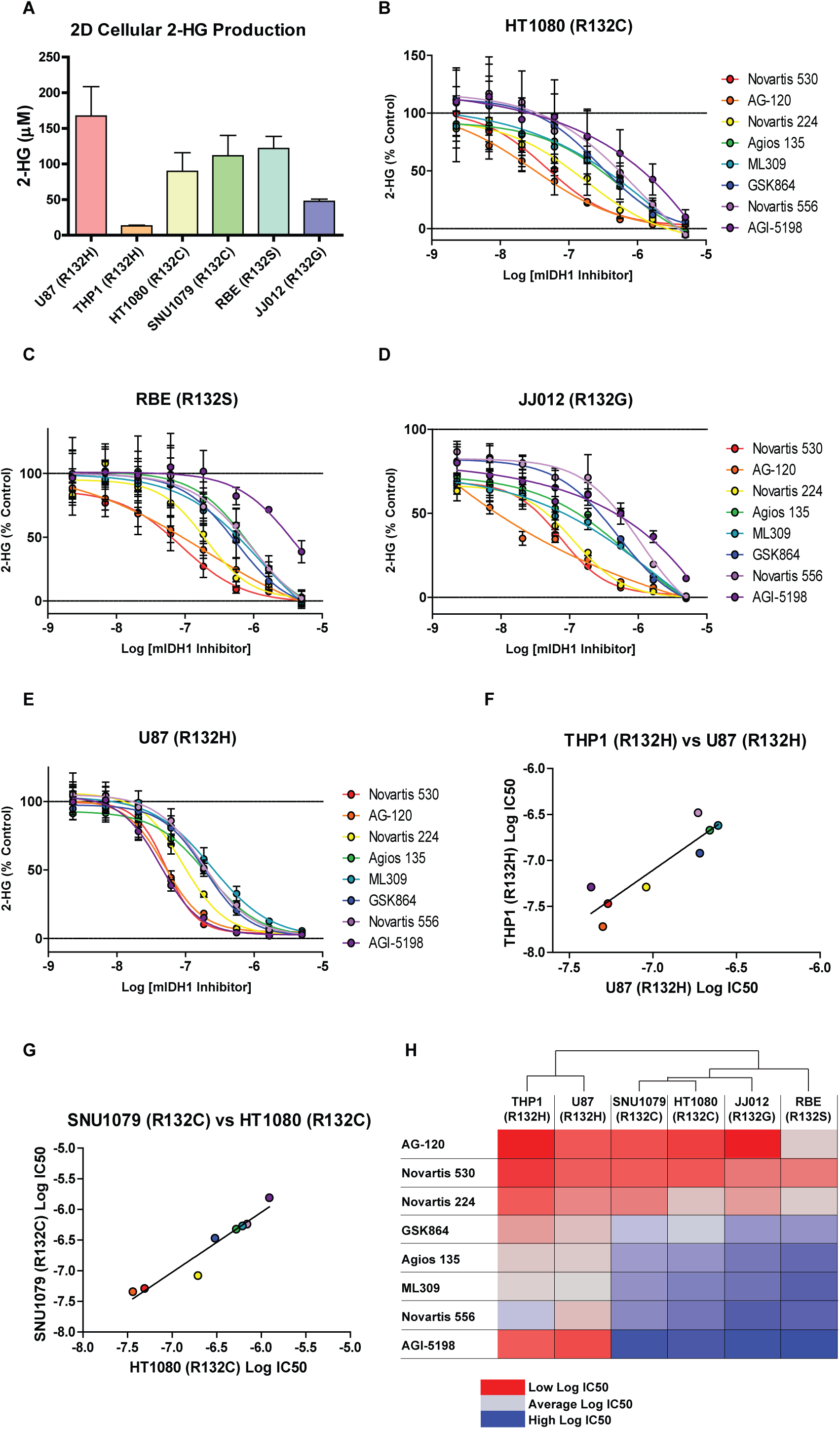
Activity of mIDH1 inhibitors in 2D cellular models. A, Average concentration of 2-HG in media produced by mIDH1 containing cell lines cultured in 2D after 48 hrs. Bars represent average and SD of 2-HG concentrations determined by mass spectrometry analysis (n=11-12). Dose-response analysis of mIDH1 inhibitor effects on cellular 2-HG production in, (B) HT1080 (R132C), (C) RBE (R132S), (D) JJ012 (R132G) (E) U87(R132H) cell lines (n=3). F, Comparison of mIDH1 inhibitor Log IC_50_ values from THP1 (R132H) and U87 (R132H) cell lines show a strong correlation (R^2^ = 0.81, P value = 0.002). G, Comparison of mIDH1 inhibitor Log IC_50_ values from SNU1079 (R132C) and HT1080 (R132C) cell lines show a strong correlation (R^2^ = 0.93, P value = 0.0001). H, Heat map of Log IC_50_ values of mIDH1 inhibitors from all six cell lines. Cell line dendrogram was generated using the UPGMA clustering method with Euclidean distance measures.

### Cellular characterization of mIDH1 inhibitors

Cellular inhibition of mIDH1 was assessed in four human cell lines derived from various malignancies known to harbor somatic IDH1 mutations - HT1080 (R132C) fibrosarcoma (33); SNU1079 (R132C) cholangiocarcinoma (34); RBE (R132S) cholangiocarcinoma (34); JJ012 (R132G) chondrosarcoma (35) - and two cell lines engineered to express mIDH1 - U87 (R132H) glioblastoma (18) and THP-1 (R132H) AML (28) (Supplementary Table 2). Inhibition of the target was assessed by quantitative measurement of 2-HG levels in extracellular media of monolayer (or 2-dimentional, 2D) cultures following incubation (48 hrs) with inhibitors. Baseline media 2-HG concentrations after 48 hours of incubation ranged from 167 μM (U87-R132H) to 13 μM (THP1-R132H) (Figure 2A). mIDH1 inhibitors were found to be active across all cell lines, demonstrating similar IC_50s_ for all cell backgrounds. Consistent with biochemical activity, the most potent inhibitors in all six cell lines (representing four different R132 mutations) were AG-120 and Novartis 530, with IC_50_ values in the range of 50-220 nM. Interestingly, Agios probe AGI-5198 reproducibly demonstrated potent, selective inhibition in the two R132H cell lines (U87 IC_50_ = 40 nM and THP-1 IC_50_ = 50 nM) compared with other cell lines (>1 μM, Figures 2B-F, Supplemental Figure 2B-C, and Supplemental Table 1).

Among the 6 cell lines tested, two harbored the R132H mutation and two others possessed the R132C mutation, allowing for a comparison of inhibitor activity against a specific R132 mutation in different cellular backgrounds. A strong correlation was observed between inhibition in THP-1 and U87 cells (R132H, R^2^ = 0.81) and between SNU1079 and HT1080 cells (R132C, R^2^ = 0.93). Hierarchical clustering of cell line sensitivity (IC_50_) further supports this strong correlation by mutation type (Figure 2H). Cell viability was also examined at 48 hrs and no significant cytotoxicity was observed for any of the mIDH inhibitors (not shown).

### Diaphorase-coupled high-throughput assay for measuring 2-HG levels

Quantitative 2-HG measurement relies primarily on mass spectrometry (MS) analysis, which is time-consuming and can be cost-prohibiting. To address this, we optimized a high-throughput method to quantify the effects of small molecule mIDH1 inhibitors on 2-HG production levels in HT1080 mIDH1 R132C fibrosarcoma cells. Previous work has shown that D-2-hydroxyglutarate dehydrogenase (D2HGDH) from *Acidaminococcus fermentans* catalyzes the oxidation of 2-HG to α-KG, as well as the reduction of the cofactor NAD+ into NADH as a product (26). By modifying reaction conditions previously developed for this bacterial enzyme, we developed a system to rapidly enable deproteination of spent cell media, and enzymatically-couple the D2HGDH reaction with diaphorase activity through NADH to provide a fast and inexpensive high-throughput fluorescence assay to quantify 2-HG production in cells (Fig. 3A-B).

**Figure 3.**
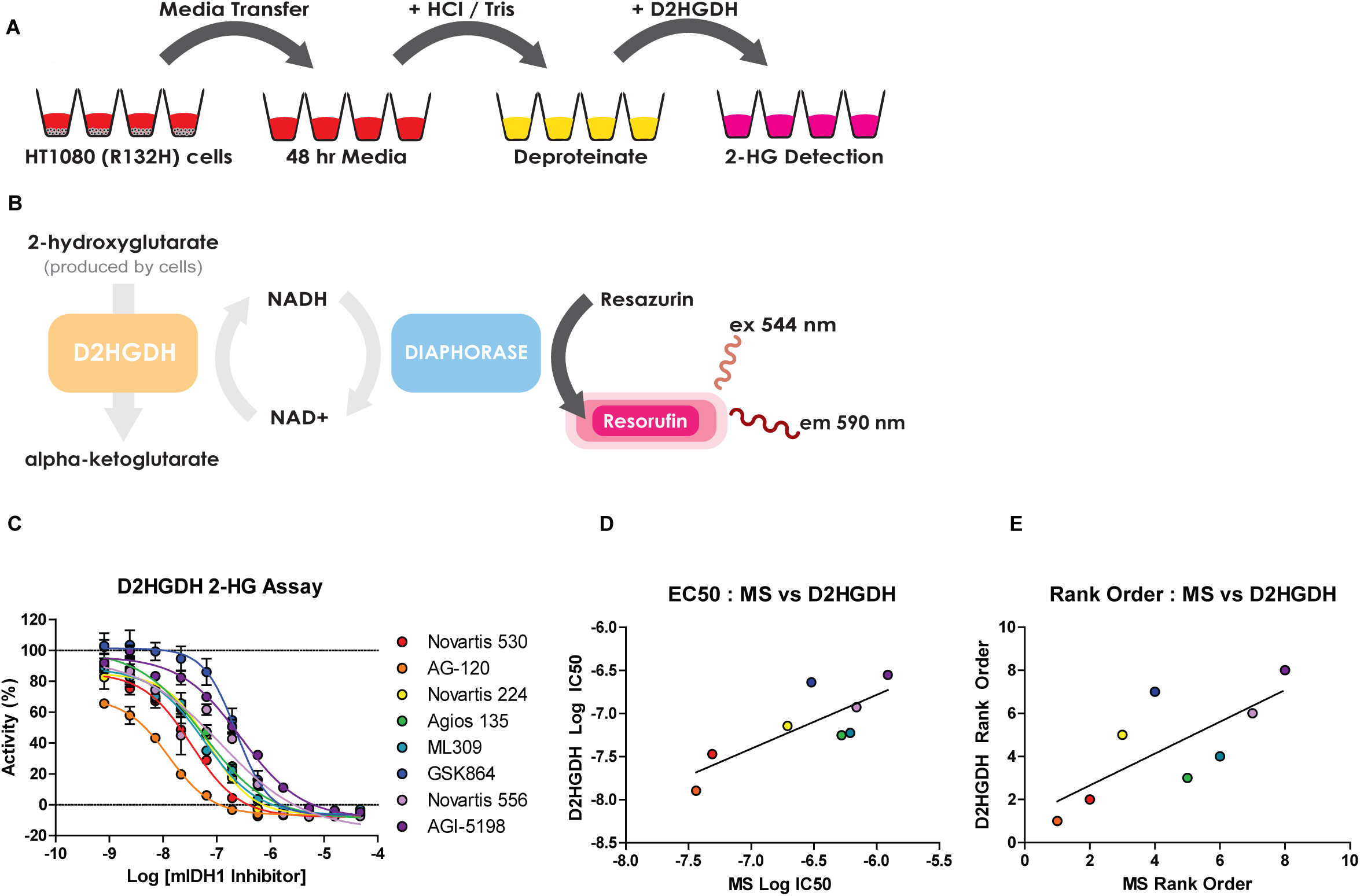
Detection of 2-HG using a fluorescent D-2-Hydroxyglutarate Dehydrogenase (D2HGDH) enzymatic assay. A-B, Scheme of the cell-based diaphorase coupled D2HGDH assay. Media was transferred to a new plate 48 hours after cells were plated and compounds were added. Following a deproteination step, D2HGDH enzyme reaction mix was added for fluorescent quantitation of 2-HG concentrations. C, Dose-response analysis of mIDH1 inhibitor effects on 2-HG production utilizing the D2HGDH assay (n=2). E, Comparison of mIDH1 inhibitor Log IC_50_ values observed from mass spectrometry analysis (MS) and the D2HGDH assay (R ^2^ = 0.63, P value = 0.019). F, Comparison of rank order based on Log IC_50_ values of mIDH1 inhibitors generated from MS analysis and the D2HGDH assay (R ^2^ = 0.55, P value = 0.036).

Detection limits and assay linearity were determined using a standard curve of 2-HG at concentration range of 8 μM to 4 nM from which a linear relationship was determined (R^2^=0.93) (Supplemental Figure 2D). Baseline 2-HG concentrations after 48 hours was 1.39 μM. Using the optimized assay conditions, we tested the inhibitors in 384-well plate format at concentrations from 47 μM to 0.8 nM. The assay demonstrated robust performance, with an average Z’ of 0.59 and an average signal to background ratio of 3.44, indicating the protocol is suitable for measuring 2-HG levels secreted from cells. AG-120 and Novartis 530 showed the greatest decrease of 2-HG levels in a dose-dependent manner, with IC_50_s of 13 nM and 34 nM, respectively (Fig. 3C). Additionally, results from the diaphorase-coupled assay correlated well with the MS-based 2-HG data, in terms of both biochemical IC_50_ values (R^2^=0.62), and overall rank order (R^2^=0.55) (Fig. 3D-E).

### Comparison of cellular target engagement of mIDH1 inhibitors

A challenging hurdle for small molecule probe/drug development is the determination of cellular target engagement. Cellular thermal shift assays (CETSA) provide a label free biophysical method that facilitates the direct measurement of cellular target engagement (24). In order to assess and compare the cellular target engagement of mIDH1 inhibitors, we utilized the engineered U87 (R132H) glioblastoma cell line. An initial cellular thermal melt analysis performed over the temperature range of 37 – 73°C found an optimal isothermal melting temperature of 59°C for mIDH1 R132H (Figure 4A-B). An isothermal dose-response (100 – 0.02 μM) of the panel of mIDH1 inhibitors was then performed at 59 °C for 3 minutes (Figure 4C-D). Target engagement resulted in stabilization of mIDH1 protein compared with DMSO (Figure 4C). Area under the curve (AUC) analysis for each dose-response curve revealed that while a majority of the inhibitors induced strong thermal stabilization of mIDH1-R132H, AG-120 and Novartis 530 are among the top compounds (Figure 4E). With respect to the other mIDH1 assays, IC_50_ values in the CETSA assay demonstrated a lack of correlation with IC_50s_ in the biochemical R132H assay (R^2^ = 0.18), while a stronger correlation was seen between CETSA and cellular (U87-R132H) mIDH1 inhibition (R^2^ = 0.54) (Figure 4F-G). Interestingly, this correlation greatly increases (R^2^ = 0.78) when AGI-5198 was excluded from the analysis.

**Figure 4.**
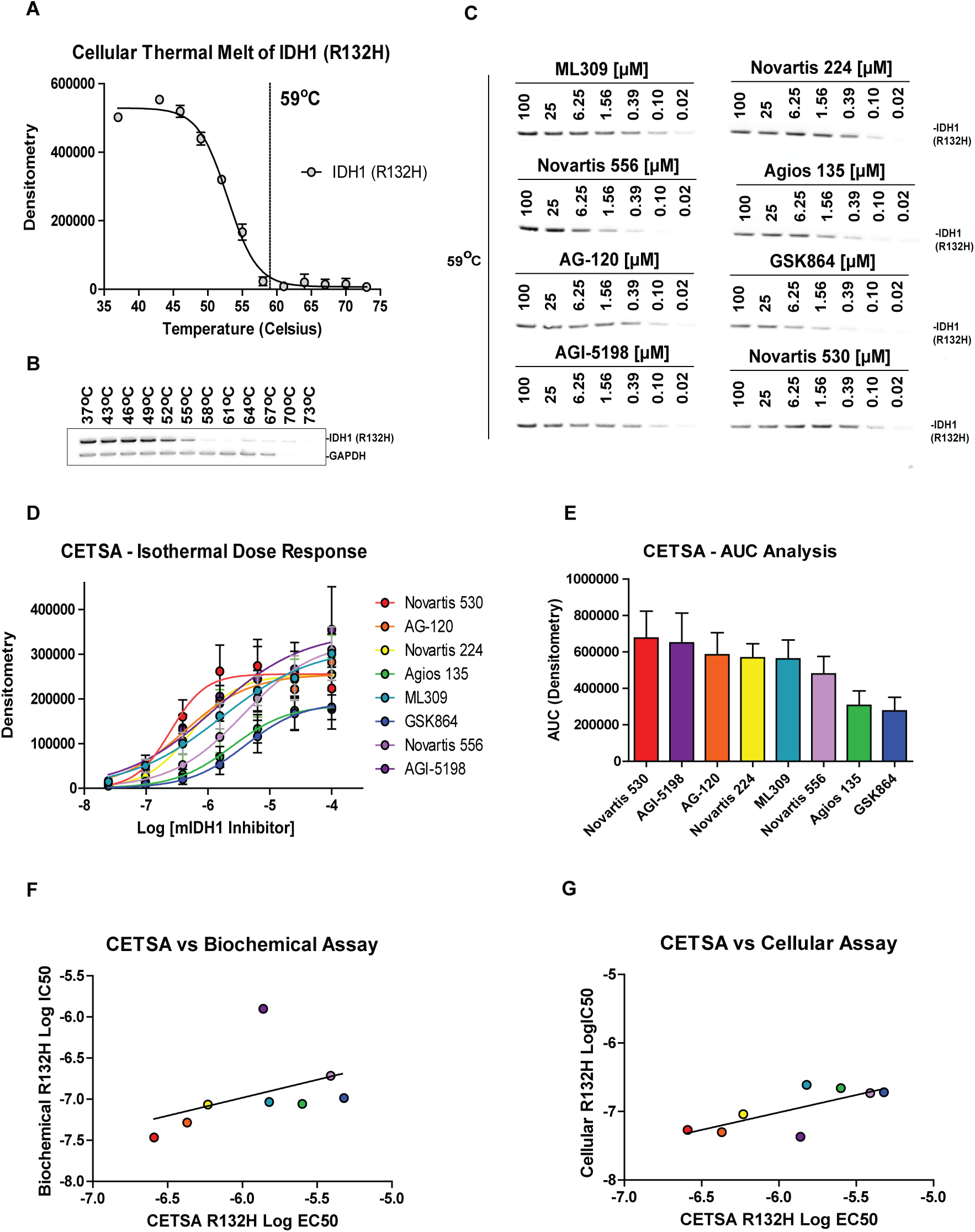
mIDH1 inhibitors increase the thermal stability of cellular mIDH1 (R132H) enzyme in the cellular thermal shift assay. A, Densitometry analysis of mIDH1 enzyme after a cellular thermal melt of U87 (R132H) cells (n=2) demonstrates 59°C as an optimal temperature for isothermal dose-response analysis. B, Representative western blot of the cellular thermal melt of U87 (R132H) cells using a temperature range of 37^°^C – 73°C. C, Represented western blot of an isothermal dose-response at melting temperature of 59°C demonstrating cellular mIDH1 enzyme thermal stabilization with mIDH1 inhibitors. D, Isothermal dose-response analysis of mIDH1 inhibitors’ stabilization of the cellular mIDH1 enzyme (n=4). E, Area under the curve analysis derived from the isothermal dose response curves. F, Comparison of mIDH1 inhibitor Log EC_50_ values from the biochemical assay (mIDH1 (R132H)) and the cellular thermal shift assay shows no correlation (R ^2^ = 0.18, P value = 0.234). G, Comparison of mIDH1 inhibitor Log EC_50_ values from the cellular assay (U87(R132H)) and the cellular thermal shift assay shows a correlation (R ^2^ = 0.54, P value = 0.038).

### mIDH1 inhibitors increase monocyte differentiation in mIDH1 expressing THP1 leukemia cells

High levels of 2-HG have been shown to repress expression of lineage-specific differentiation genes, block cell differentiation, and correlates with an increase in repressive histone methylation marks (36). Inhibitors of mIDH enzymes have been shown to induce differentiation of mIDH bearing AML cell lines and primary patient cells (21). In order to compare the ability of mIDH1 inhibitors to induce differentiation of AML cells, we implemented a cell differentiation assay using human THP1 leukemia cells engineered to express either wild type IDH1 (THP1-WT) or mIDH1-R132H (THP1-R132H) under a doxycycline-inducible promoter (28,29). Mutant or wild type IDH1 gene expression was induced with doxycycline (0.5 μM) for four days prior to the induction of monocytic differentiation with phorbol-12-myristate-13-acetate (PMA) (50nM), ultimately resulting in cellular adherence (Figure 5A shows a schematic representation of the experiment and results). Differentiation was determined by imaging and quantifying the number of cells that adhered to each well in a 96-well plate (Figure 5B-C). After treatment with PMA, THP1 (R132H) cells were found to have reduced levels of adherent cells compared to THP1 (WT), indicating that high 2-HG levels may block cell differentiation (Figure 5C). Treatment of the THP1 (R132H) cell line with 0.5 μM of mIDH1 inhibitors revealed an increase in the number of adherent cells compared to DMSO alone (Figure 5D and Supplemental Figure 3). Additionally, co-treatment of THP1 (R132H) cells with the cell-permeable octyl-2-HG and mIDH1 inhibitors, specifically Novartis 530 and AG-120, greatly diminished the number of differentiated cells compared to mIDH1 inhibitor treatment alone, confirming elevated 2-HG levels are capable of blocking THP1 differentiation (Figure 5E).

**Figure 5.**
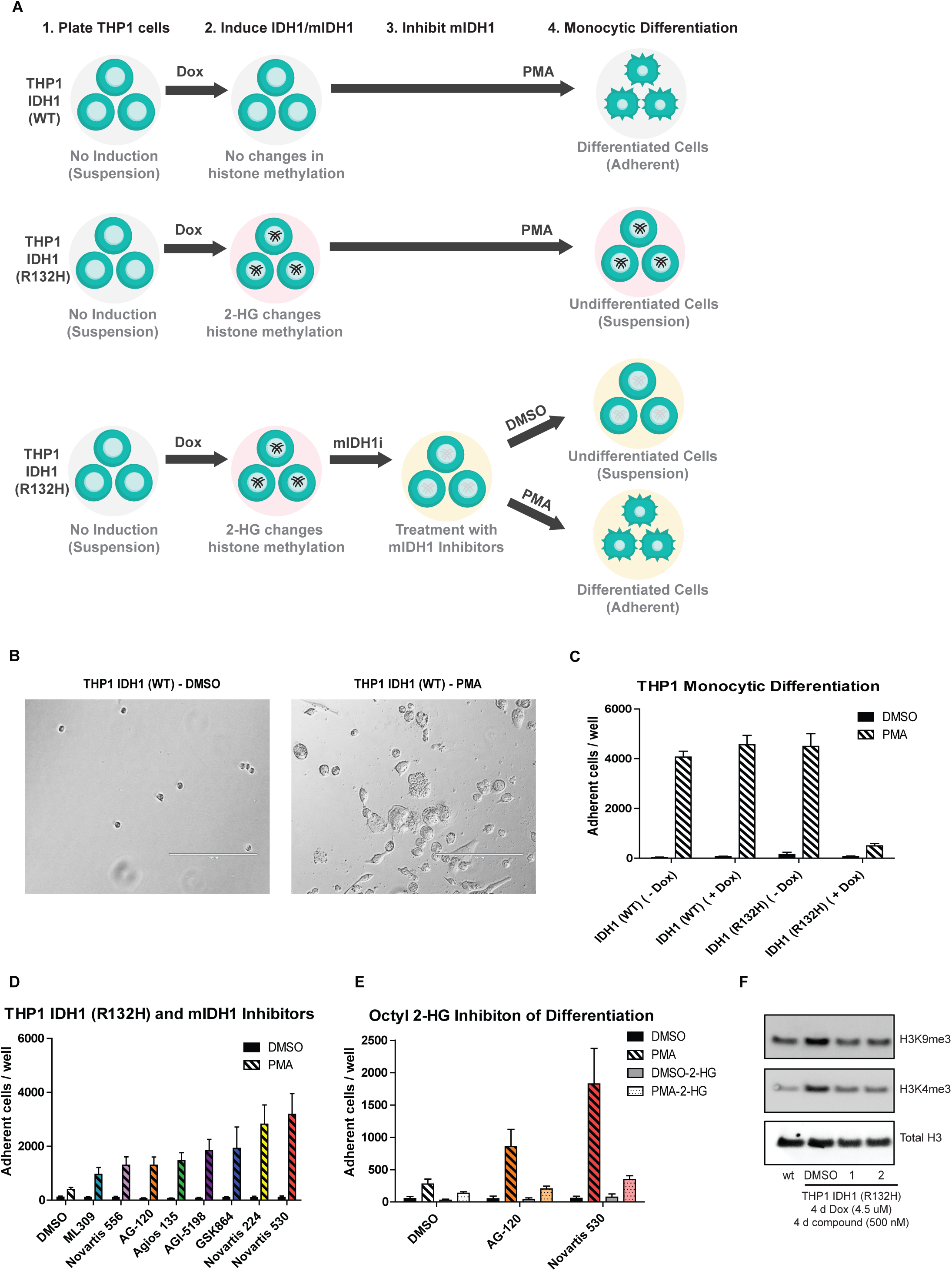
mIDH1 inhibitors rescue monocytic differentiation and histone methylation of mIDH1 expressing THP1 leukemic cells. A, Schematic representation of the differentiation assay. Engineered THP1 cells grown in suspension were treated with doxycycline to induce WT IDH1 (top) or mIDH1-R132H (middle) expression followed by PMA to induce differentiation. Induction of mIDH1 increases 2-HG levels, alters histone methylation, and blocks differentiation. mIDH1 inhibitors block mIDH1 activity and reduce 2-HG (bottom), allowing for the rescue of PMA induced monocytic differentiation. B, Representative images of adherent doxycycline induced THP1 IDH1 (WT) cells treated for 7 days with DMSO (left) or PMA (right) (scale = 200 μm). C, Number of adherent THP1 leukemic cells expressing either WT or mIDH1 R132H after 7 days of treatment with DMSO or 50 nM PMA. Bars represent average and SD of number of adhered cells / well (n=16). D, Number of adherent mIDH1 expressing THP1 leukemic cells after 5 days of treatment with either DMSO or 50 nM PMA following 4 days of treatment with 500 nM of mIDH1 inhibitors. Bars represent average and SD of number of adhered cells / well (n=16). E, Number of adherent mIDH1 expressing THP1 leukemic cells after 5 days of treatment with either DMSO or 50 nM PMA following 4 days of treatment with 50 μM mIDH1 inhibitors in the presence or absence of 300 μM octyl-2-HG. F, Representative western blot from purified histones showing changes in H3K9me3 and H3K4me3 methylation levels in wild type IDH1 and mIDH1 expressing THP1 leukemic cells after 4 days of doxycycline following treatment with either vehicle, AG-120 (1), or Novartis 530 (2) (n=2).

Given that excess 2-HG levels have been shown to increase methylation through inhibition of cellular demethylases, histone tri-methylation levels were examined in treated and untreated THP1 cells. H3K9me3 and H3K4me3 levels were found to be increased in THP1 (R132H) cells compared to THP1 (WT) cells. Furthermore, treatment of THP1 (R132H) cells with either Novartis 530 or AG-120 reduced these elevated levels of tri-methylation (Figure 5F).

### mIDH1 inhibitors reduce 2-HG levels in spheroid models

To evaluate the effect of mIDH1 inhibitors in more relevant models of solid tumors, we adapted a 3-dimensional (3D) multicellular tumor spheroid assay to monitor spheroid formation and 2-HG production. Small molecules can have diminished efficacy in multicellular tumor spheroid models due to limited penetration and the heterogeneous nature of the 3D tumor spheroid (37,38), but no reports of the activity of mIDH1 inhibitors in 3D models exist. We began by testing the 6 aforementioned cell lines for their ability to form spheroids in ultra-low attachment plates at varying cell concentrations (1000 – 5000 cells/well). Four cell lines (HT1080 fibrosarcoma, JJ012 chondrosarcoma, U87 glioblastoma and RBE cholangiocarcinoma), each with a different R132 mutation, were shown to form spheroids (Fig 6A). Overall, spheroids produced less extracellular 2-HG than cells cultured in monolayer (Fig 6B).

**Figure 6.**
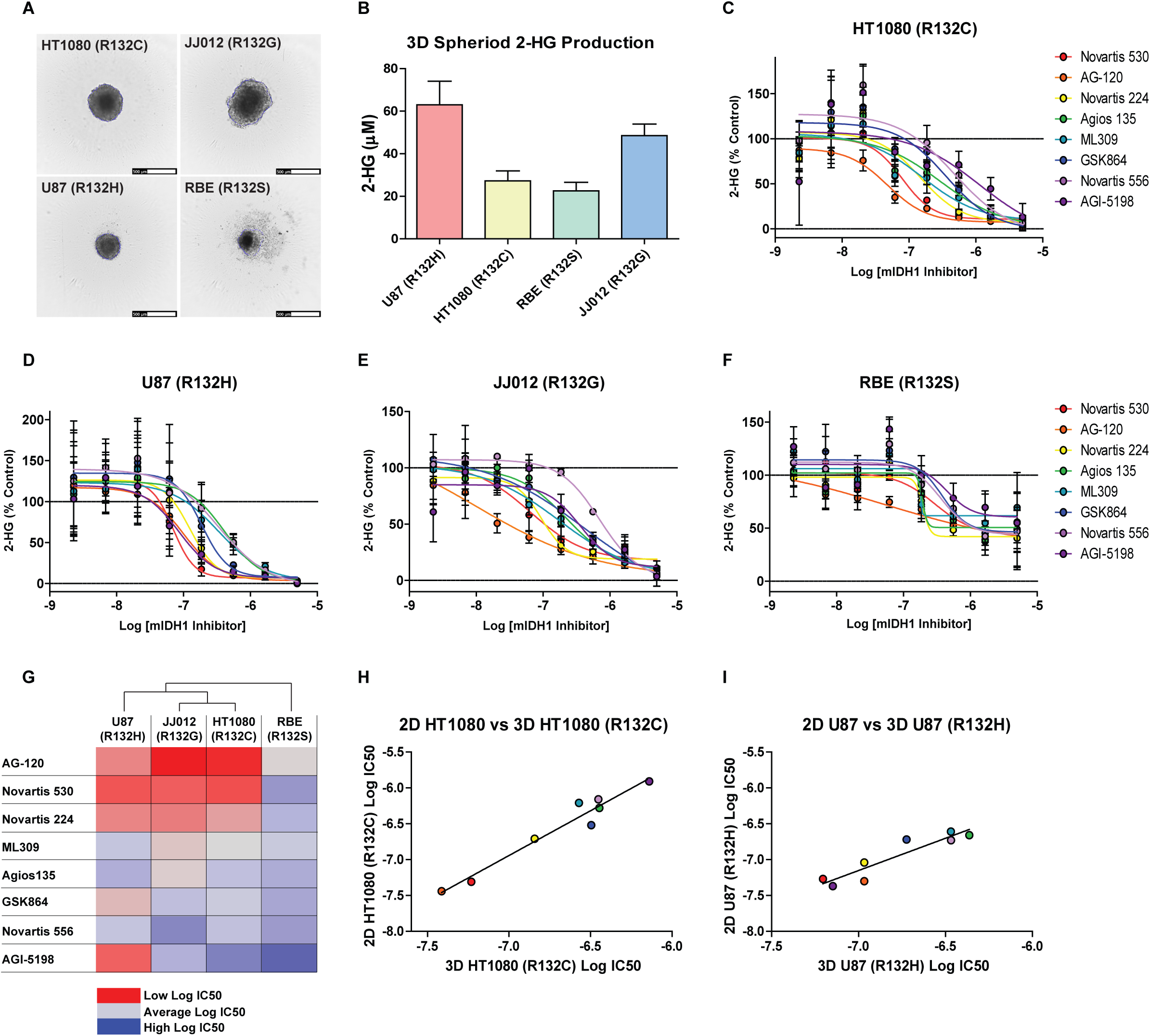
Activity of mIDH1 inhibitors in 3D cellular spheroids. A, Representative images of 3D cellular spheroids of HT1080 (top left), JJ012 (top right), U87 (R132H) (bottom left), and RBE (bottom right) cells (scale = 500 μm). B, Concentration of 2-HG in media produced from 3D cellular spheroid cultures after 4 days. Bars represent average and SD of 2-HG concentrations determined by MS analysis (n=12). Dose-response analysis of mIDH1 inhibitor effects on 2-HG production in (C) HT1080(R132C), (D) U87(R132H), (E) JJ012(R132G), (D) RBE(R132S) 3D cellular spheroids (n=3). G, Heat map of Log IC_50_ values of mIDH1 inhibitors from the four 3D cellular spheroids. Cell line dendrogram was generated using the UPGMA clustering method with Euclidean distance measures. H, Comparison of mIDH1 inhibitor Log IC_50_ values between 2D and 3D HT1080 (R132C) cellular assays show a strong correlation (R ^2^ = 0.95, P value = <0.0001). I, Comparison of mIDH1 inhibitor Log IC_50_ values between 2D and 3D U87 (R132H) cellular assays show a strong correlation (R ^2^ = 0.86, P value = 0.0009).

Using these simple 3D models of solid tumors, we assessed the ability of inhibitors to reduce 2-HG production. All mIDH1 inhibitors tested demonstrated complete suppression of 2-HG against all cell lines except the RBE cells (Fig 6C-G), which had reduced efficacy and no effect on rank-order of potency. Interestingly, AG-120 and Novartis 530 were the two most efficacious inhibitors in both monolayers (2D) and 3D cultures (Figure 6G-I). Moreover, the activity of inhibitors in 2D and 3D models demonstrated strong positive correlations, particularly in the HT1080 (R^2^ = 0.95) and U87 (R^2^ = 0.86) cell lines (Figure 6H-I).

### *In vitro* drug metabolism and pharmacokinetic properties of mIDH inhibitors

Drug metabolism and pharmacokinetic properties of probe compounds and drug candidates ultimately determine how these compounds behave/perform in preclinical animal models and clinical trials (39). Using robust, automated, high-throughput *in vitro* assays, we measured the solubility, permeability, metabolic stability, and plasma protein binding properties for the panel of mIDH1 inhibitors. Aqueous solubility of the mIDH1 inhibitors was assessed using a kinetic solubility determination in an aqueous solution at pH 7.4 (40). While a wide range of solubilities were observed (<0.1-147.9 μM), only Agios 135 and GSK864 had solubility concentrations that approached their respective IC_50_ values from the 2-HG cell-based assay (Supplemental Table 1). On the other hand, Novartis 530 and AG-120 had the greatest separation between solubility and IC_50_ values. Parallel artificial membrane permeability assays (PAMPA) were used to model passive, transcellular permeability across the gastrointestinal tract (GIT) (41). All compounds tested were characterized as highly permeable (>100 × 10^−6^ cm/s) (Supplemental Table 1). Hepatic microsomal stability half-life (t_1/2_) values in three species (mouse, rat and human) were determined using a multiple time-point metabolic stability assay (42). AG-120 had the highest metabolic stability, while the rest of the mIDH1 inhibitors were classified as low stability compounds with t_1/2_ less than 10 min (Supplemental Table 1). Human plasma protein binding revealed a very high (>96%) protein binding for most mIDH1 inhibitors, while AG-120 and Novartis 556 demonstrated the greatest unbound fractions (∼ 10% unbound) (Supplemental Table 1). Caco-2 permeability was measured to gain insight into potential interactions with membrane transporters. While Agios 135 and AGI-5198 had increased efflux ratios suggesting potential interactions with membrane transporters, AG-120 demonstrated the highest efflux ratio at 56 with very low apical to basal transport (Supplemental Table 1).

The apparent basal to apical transport of AG-120 in Caco-2 cells was suggestive of an active drug transporter such as the multidrug transporter P-glycoprotein (P-gp, MDR1, *ABCB1*) (43). AG-120 is in human clinical trials, where P-glycoprotein at the blood-brain barrier (BBB) might limit drug uptake in the CNS (44). To investigate this, transwell transport of AG-120 was also assessed in MDCK cells transfected with P-gp. AG-120 demonstrated strong basolateral to apical transport and almost no apical to basolateral transport. This imbalance was completely reversed in the presence of the P-gp small molecule inhibitors tariquidar and verapamil (Supplemental Figure 5A). For comparison, AG-120 demonstrated a greater efflux ratio (AG-120 efflux ratio >350) than the avid P-gp substrate loperamide (efflux ratio ∼350), which cannot cross the BBB (Supplemental Figure 5B) (45).

*In vitro* predictive toxicity assays were also utilized to determine if mIDH1 inhibitors may cause cytotoxicity in human hepatoyctes or inhibit hERG channels (46). All mIDH1 inhibitors show little to no cytotoxicity after 72 hr incubation with human hepatocytes, except at the highest doses tested (46 μM) (Supplemental Figure 5C). Activity at the well-known anti-target, the hERG channel, was assessed using the hERG FluxOR-based thallium flux assay. All compounds tested had IC_50_ values above 10 μM except for GSK864 (IC_50_ = 4.7 μM) (Supplemental Figure 5D).

## Discussion

There are multiple mIDH1/2 inhibitors currently in clinical trials (47). Agios has made the most progress, advancing its mIDH2 inhibitor, AG-221 (Enasidenib), into a phase III open-label clinical trial in 14 countries for late stage AML harboring an IDH2 mutation (NCT02577406). Agios has also advanced its mIDH1 inhibitor, AG-120 (Ivosidenib), into a phase III clinical trial for metastatic cholangiocarcinoma (NCT02989857). Additionally, Agios is pursuing a brain-penetrant dual mIDH1/2 inhibitor, AG-881, currently in phase I for advanced solid tumors including gliomas containing an IDH1/2 mutation (NCT02481154). Bayer (BAY1436032), Novartis (IDH305), and Forma Therapeutics (FT-2102) all have mIDH1 inhibitors currently in phase I clinical trials for either solid tumors or refractory AML (NCT02746081, NCT02381886, and NCT02719574). Recently, Bayer disclosed the structure of their clinical mIDH1 inhibitor BAY 1436032, showing favorable oral pharmacokinetic properties, reasonable brain barrier penetration, and a direct interaction with the mutational residue His132 of the mutant IDH1 R132H enzyme (48). Novartis also recently published an optimized mutant IDH1 inhibitor (IDH889) similar to the chemotype studied here, with favorable oral pharmacokinetic properties and brain barrier penetration (49). Several inhibitor classes were not available to us at the time this study was undertaken, including the Bayer (BAY1436032), Forma (FT-2102), and Novartis (IDH305 and IDH889) inhibitors previously mentioned.

This assessment of multiple mIDH inhibitors across many *in vitro* assay platforms characterizes the comparative activities of these mIHD1/2 inhibitors. AG-120 and Novartis 530 rank the highest when observing biochemical potencies, reduction of 2-HG in all six cell lines, and when measuring isothermal dose response using CETSA.

Novartis 530 and 224 show the best activity in reversing 2-HG dependent differentiation of THP1 leukemic cells, while AG-120’s activity is much weaker in this assay. AG-120 has the best *in vitro* physicochemical and drug metabolism profile of all the compounds tested, yet it is a substrate for P-gp transporters preventing its passage through the BBB. This study identifies appropriate inhibitors for studying mIDH1/2 enzymes and their role in the pathophysiology of certain diseases. It also provides an opportunity to confirm that the biochemical and cell-based models utilized in previous studies are comparable. This is not necessarily clear from the literature, where individual lines have been used when reporting specific inhibitor activities. ML309 and initial Agios probes were characterized biochemically using mIDH1 R132H and mR132C homodimer protein, and in U87 (R132H) cells and HT1080 cells (18). HT1080 cells were one of the first cell lines reported to carry a somatic IDH1 mutation (33). Even though the prevalence of IDH mutations are very high in gliomas, mIDH1 or mIDH2 low-grade glioma cell lines have been difficult to generate (the TS603 oligodendroglioma harbors an IDH1 R132H mutation and grows as neurospheres rather than monolayer) (1,16), leading to the use of a variety of transfected GBM cell lines. AGI-5198 activity was reported against mutant/wild type IDH1 heterodimer protein and the aforementioned TS603 cell line (16). The GSK321 inhibitor was assessed against three mutant forms of protein (R132H, C and G), and in HT1080 cells (21). The Sanofi inhibitor was assessed against a range of mIDH1 homo- and hetero-dimer protein, and in HEK293 cells transfected with mIDH1 (R132H)(22). The mIDH2 inhibitor AGI-6780 was assessed in HEK cells transfected with R140Q IDH2 (23). Here, we report the first side-by-side comparison of a diverse set of mIDH1 inhibitors across multiple *in vitro* assay platforms.

Comparative efficacy of multiple chemotypes against an individual target is not commonly undertaken, but given the strong interest in mIDH as an oncotarget and rapid emergence of multiple parallel inhibitor development programs, we were interested in how these chemotypes compared in assays that reported on target activity from biochemical to cellular. A side-by-side comparison of available small molecules against multiple cell lines may appear to be designed to identify the ‘best’ inhibitor. However, it is important to comprehensively evaluate these compounds to better understand their respective limitations; the availability of multiple mIDH1 and mIDH2 inhibitors will help ensure that observed phenotypes for individual compounds are truly on-target. The majority of inhibitors demonstrated activity in line with published reports, though in our hands the Sanofi inhibitor was not as efficacious. The AG-120 and Novartis 530 were the most active across a majority of assay platforms.

We found that inhibition across multiple IDH1 mutations was generally consistent, and that this translated to cellular inhibition of mIDH1, both in engineered and somatic cell lines. In addition, we miniaturized a D2HGDH florescence-based assay to determine 2-HG concentrations in cell medium. This assay provides an accessible platform for research laboratories to measure 2-HG without the need for quantitative MS. We utilized the assay developed by Jorg Balss and colleagues that coupled the D2HGDH enzyme to resazurin to measure intracellular and extracellular 2-HG (26). Adnan Elhammali and colleagues subsequently reported an HTS screen using a similar approach in which they showed that Zaprinast reduces 2-HG by perturbing glutamine metabolism, although the HTS assay utilized *E. coli* PHGDH, whose primary substrate is 3-phosphoglycerate limiting its utility as a specific reporter assay (50). The D2HGDH-coupled assay used here was cost effective, readily scaled down and semi-automated with liquid-handling, providing a high-throughput convenient method to quickly analyze 2-HG containing media samples from cells.

CETSA is a relatively new assay used to interrogate target engagement independent from function (51), and we found it to be generally informative of mIDH1 engagement and correlated to cellular inhibition of mIDH1. Given the opportunity to use 2-HG as a biomarker *in vivo* (along with MRI modalities), mIDH1 may provide a strong opportunity to correlate xenograft target engagement using CETSA with target inhibition. The development of 3D tumor spheroids generated from cell lines (reported here) demonstrated that 2-HG production is measureable, and inhibitor efficacy was retained in solid tumor models and correlated with activity in monolayer cells. Interestingly, the greatest divergence from ranked activity was observed in the differentiation assay where AG-120 had reduced efficacy in rescuing the differentiation phenotype. While AG-120 demonstrated superior physicochemical and drug metabolism properties, its susceptibility to transport by P-gp may limit activity in drug-resistant mIDH1 cells, and may restrict brain penetrance (52). Our panel of assays, some of which are novel and amenable for high-throughput, provide a comprehensive platform for drug discovery campaigns to identify additional mIDH1 inhibitors.

## Acknowledgement

This work was supported by the National Cancer Institute’s (NCI) Experimental Therapeutics Chemical Biology Consortium (CBC).

